# Paternal starvation affects metabolic gene expression during zebrafish offspring development and life-long fitness

**DOI:** 10.1101/2023.09.22.557632

**Authors:** Ada Jimenez-Gonzalez, Federico Ansaloni, Constance Nebendahl, Ghazal Alavioon, David Murray, Weronika Robak, Remo Sanges, Ferenc Müller, Simone Immler

**Affiliations:** Institute of Cancer and Genomic Sciences, College of Medical and Dental Sciences, University of Birmingham, Vincent Drive, Edgbaston, B15 2TT, Birmingham, UK; Department of Evolutionary Biology, Uppsala University, Uppsala, Sweden; Central RNA Laboratory, Istituto Italiano di Tecnologia (IIT), Genova, Italy; Area of Neuroscience, Scuola Internazionale Superiore di Studi Avanzati (SISSA), Trieste, Italy; School of Biological Sciences, University of East Anglia, Norwich, United Kingdom; Centre for Environment, Fisheries, and Aquaculture Science, Pakefield Road, Lowestoft, NR33 0HT, UK

**Keywords:** starvation, paternal, non-genetic, development, transcriptome

## Abstract

Dietary restriction is a putative key to a healthier and longer life, but these benefits may come at a trade-off with reproductive fitness and may affect the following generation(s). The potential inter- and transgenerational effects of starvation are particularly poorly understood in vertebrates when they originate from the paternal line. We utilised the externally fertilising zebrafish amenable to a split-egg clutch design to explore the male-specific effects of starvation on fertility and fitness of offspring independently of maternal contribution. Eighteen days of fasting resulted in reduced fertility in exposed males. While average offspring survival was not affected, we detected higher larval growth in offspring from starved males and increased malformation rates at 24 hours post fertilisation in the F2 embryos produced by the offspring of the starved males. The transcriptome analysis of embryos from starved and fed fathers revealed robust and reproducible induction of muscle composition genes and a contrasting repressive effect on lipid metabolism and lysosome genes. A large proportion of these genes showed enrichment in the yolk syncytial layer suggesting gene regulatory responses associated with metabolism of nutrients through paternal impact on extra embryonic tissues which are loaded with maternally deposited factors. We compared the embryo transcriptome to adult transcriptome datasets and demonstrated comparable repressive effects on metabolism-associated genes. These similarities suggest a physiologically relevant, directed and potentially adaptive response transmitted by the father, independently from the offspring’s nutritional state, which was defined by the mother.

## Introduction

Dietary restriction in the form of caloric restriction and intermittent fasting is a putative key to the extension of lifespan and health-span (Adler & Bonduriansky, 2014; Kapahi et al., 2017). The benefits of intermittent fasting for organismal health have been reported in a range of invertebrates and vertebrates including humans (Honjoh et al., 2017; Mitchell et al., 2019; Pan et al., 2022). Despite its beneficial effects on organismal health, dietary restriction may have side effects and affect for example reproductive fitness, although the effects reported so far vary. In females, dietary restriction appears to generally have positive effects on egg quality and extend the reproductive capacity (Nehra et al., 2012; Sun et al., 2021). The effects of diet on male reproduction are less clear. Male dietary regimes can alter sperm quality and fertility and high-calory diets generally have negative effects on male fertility (e.g. Rato et al. 2014 (Rato et al., 2014)) whereas the effects of dietary restriction on male fertility and reproduction vary across species and studies. Dietary restriction had a negative effect on sperm count and sperm motility and capacitation ability in young two-months old Sprague Dawley rats (Li et al., 2021). However, it had no effects on 2.5-months old Sprague Dawley rats that were also exposed to heat stress (Aydilek et al., 2015) or led to a positive effect in aged 12-months old Wistar rats (Jesús et al., 2022). A randomised trial of dietary restriction in obese men showed an increase in sperm count and sperm concentration in the trial group under caloric restriction (Andersen et al., 2022). These conflicting results on the effects of dietary restriction on reproductive fitness suggest species and context dependence and warrant further investigation.

In addition, it is unclear how parental dietary restriction affects future generations. In the nematode *Caenorhabditis elegans*, dietary restriction results in lifespan expansion in the individuals undergoing the treatment with extended effects on up to three generations (Ivimey-Cook et al., 2021). Similarly, in human populations, famine experienced by one generation may lead to increased risk of hyperglycemia for at least two subsequent generations (Li et al., 2017). However, in these studies, both male and female gametes were produced under the conditions of dietary restriction, or the exposures occurred prenatally and hence it is unclear how important paternal adult fasting is for offspring fitness.

The modulation of the physiological responses upon exposure to dietary restriction occurs through the alterations of key metabolic pathways and gene expression. This includes changes in well conserved nutrient sensing pathways like AMPK or the mechanistic target of rapamycin (mTOR) which is inhibited by caloric restriction and may trigger autophagy, favouring the recycling of cellular components and impeding cell growth (Efeyan et al., 2013). These changes are accompanied by alterations of several other pathways because of the metabolic switch experienced by the cells. Salem et al. (2007) (Salem et al., 2007) found that food deprivation induced changes in expression of glucose, lipid metabolism, blood function and immune response-related genes in the liver of rainbow trout *Oncorhynchus mykiss*. Lipid and insulin metabolism related genes are also deregulated in a model of skeletal muscle aging in rats exposed to caloric restriction between 15 and 30 months of age (Ham et al., 2022). However, whether and how these changes in nutrient sensing and metabolic rates in starved individuals may reflect potential lasting impact that is transmitted to the next generation is little understood. Here, we explicitly explore the effects of dietary restriction by short-term starvation on male reproductive fitness and offspring performance and investigate the underlying molecular profiles using the zebrafish *Danio rerio*.

## Materials and Methods

### Animal model

For these experiments, we used wild-type Zebrafish *Danio rerio* from the AB strain obtained from ZIRC (Zebrafish International Resource Center, University of Oregon, Eugene, USA) and maintained at the SciLifeLab zebrafish platform at Uppsala University (http://www.scilifelab.se/facilities/zebrafish/), the Controlled Ecology Facility (CEF) at the University of East Anglia (UEA, UK) and the Biomedical services unit (BMSU) fish facility at the University of Birmingham (UOB, UK) . The fish were kept in 3-l tanks in a recirculating rack system (Aquatic Habitats, Beverly, MA, USA, Z-Hab System) at 26.4 ± 1.4°C and a 12:12 diurnal light cycle.

### Feeding regime

Fish were fed three times a day with a mixture of dry pellets and live artemia. For the starvation experiments, 10-12 males were randomly split into two tanks assigned to the two experimental groups, named Control and Starved (Figure 1A). Each male was assigned an ID number and weighed and imaged for posterior identification through pigmentation and fin shape features. Males were maintained with wild-type companion females at a total fish density of 10 fish per tank for the duration of the experiment.

**Figure 1.**
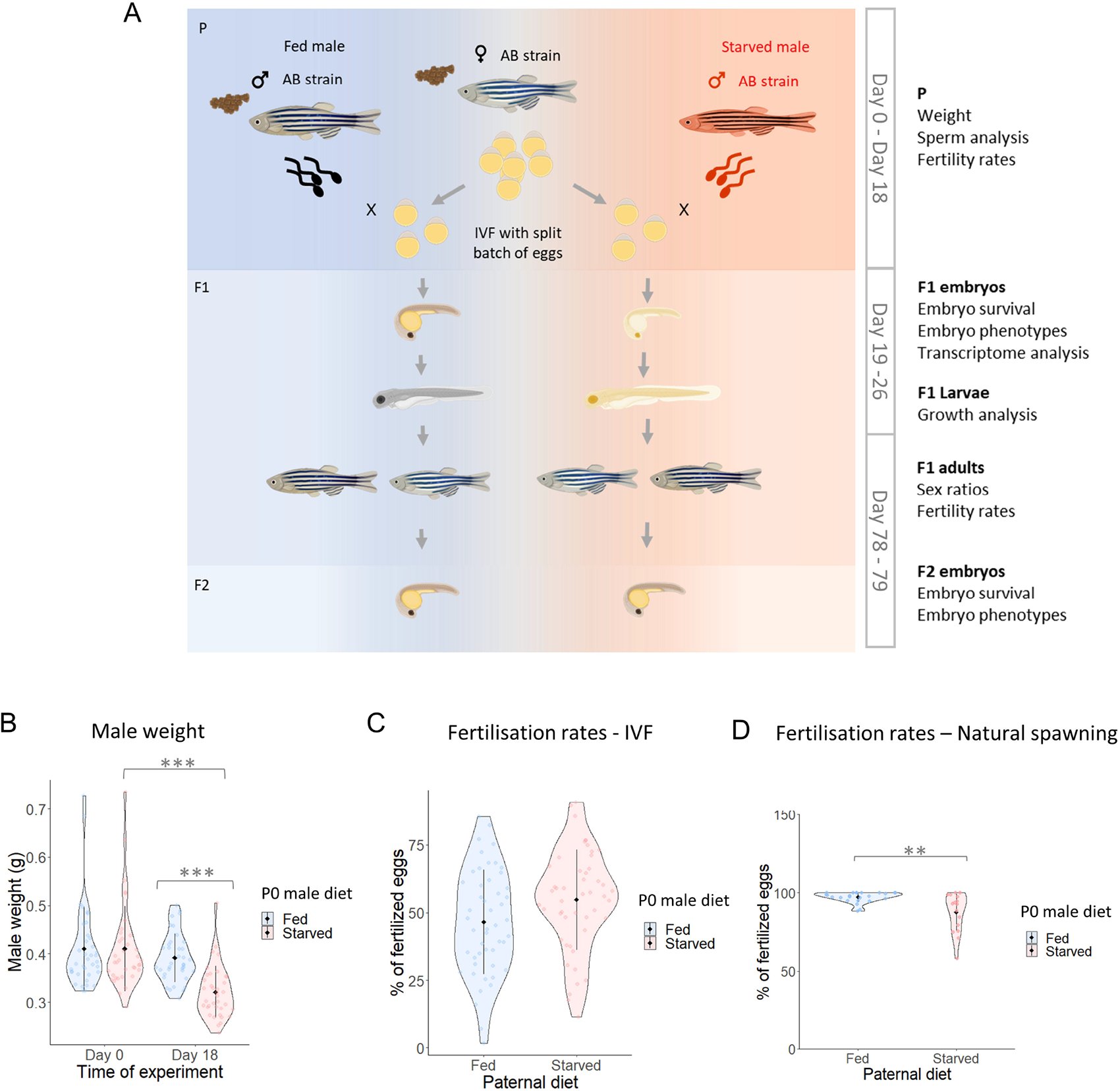
A zebrafish model to study the intergenerational effects of paternal starvation. **A.** Experimental design for IVF using a split-clutch design: All males were weighed at the beginning of the experiment (Day 0) and then randomly split into fed and starved groups. Starved males were fully deprived of food whereas fed males were fed a standard diet of a mix of dry and live (artemia) food three times a day. All males were kept with an equal number of females for the duration of the experiment. After 18 days, males were weighed again and ejaculates were collected. Eggs were collected from healthy wild-type non-experimental females and split into two equal halves for IVF. Sperm from one fed and one starved male, respectively, were used to fertilise one half of the clutch of eggs (crosses were performed with AB fish). On day 19, embryos were collected at prim-5 stage (24 hpf) for transcriptome analyses. Larval length was measured on days 5 and 8 post-fertilisation. The timeline of the experimental setup and data collected is shown on the right side of the design. **B.** Weight of fed and starved males on days O and 18 of the experiment. On day 18, the weight of starved males was significantly reduced by 21.83% in 3 independent experiments (starved group Day 0: 0.410±0.08, Day 18: 0.321±0.05, mean± SD). There was no significant difference in fed males between day O and 18. Individual data points are represented as dots within the violin plots. N=37. **C.** Percentage of fertilised eggs at 64-cell stage (2 hpf). Individual data points within the violin plot represent the average egg survival rate per male and across 7 independent IVFs (Fed males: 1777 fertilised eggs out of 3874 total eggs. Starved males: 2038 fertilised eggs out of 3947 total eggs. 4 males were used in each IVF. Sperm collected was used to fertilised eggs from 2 different females when possible). **D.** Percentage of fertilised eggs at 64-cell stage (2 hpf) and produced by natural spawning. Individual data points within the violin plots represent the average egg survival rate per male and across 3 independent rounds of breeding (Fed males: 2688 fertilised eggs out of 2784 total eggs. 6-13 males were used in 3 experiments. Starved males: 1419 fertilised eggs out of 1609 total eggs. 6-13 males were used in 3 experiments. Sperm collected was used to fertilised eggs from 2 different females when possible.).*** p < 0.001; ** p <0.01. Black bars represent the mean± SD.

Upon splitting into experimental groups, fish In the Starvation treatment were completely deprived of food while fish from the Control treatment were kept in the *ad libitum* feeding regime where they were fed three times a day as under standard lab conditions. The experiment lasted for 18 days (Fig. 1A).

### *In vitro* fertilisation

On day 18, fish were put into breeding tanks and separated from wild-type females with dividers, allowing visual and olfactory contact. Breeding tanks were then covered with black cloths to avoid any light-induced oviposition the following morning.

Females and males were prepared for *in vitro* fertilisation (IVF) and anaesthetised using 1.0–3.0 mg/l metomidate hydrochloride (AquacalmTM). Males were weighed to control for the effects of feeding regime and imaged to match their ID with the data collected on day 0. Males were then placed on a soft and wet sponge and squeezed gently in cranio-caudal direction to collect the ejaculate under a dissecting microscope (Nikon SMZ800). From each male, 0.2–0.8 μl of ejaculate were collected and transferred into a 0.2 ml Eppendorf tube containing 80 μl of Hank’s buffer (HBSS) and kept on ice for 5–10 min until IVF. Females were placed on 15 cm Petri dishes and gently stripped to obtain eggs. Clutches used for IVFs contained 20–300 high quality eggs and they were used within one minute after stripping. To create the first generation (F1), we used a split clutch design to perform IVFs (Fig. 1A). Sperm samples were mixed gently, and each ejaculate and egg clutch were divided into two parts. *In vitro* fertilisation was performed simultaneously for both experimental groups.

### Natural spawning

Reproductive fitness for experimental males (P) and male and female offspring (F1) was assessed by crossing experimental fish with non-experimental wild-type fish from an independent AB population. Some offspring (F1) from Control and Starvation males were reared into adulthood for assessment of reproductive fitness. F1 fish were kept in mixed sex groups in 3l tanks with standardised densities according to age. Larvae were kept in groups of 50 and at the age of two months, F1 fish were re-distributed into 3l tanks in mixed sex groups of 14-16 fish per 3l tank.

To assess reproductive fitness, F1 males and females were set up for natural spawning in pairs with non-experimental wildtype fish in individual breeding tanks. One day before natural spawning, male and female offspring from Control and Starvation males and wild-type males and females respectively were placed in pairs in breeding tanks where male and female were separated by a divisor. Allowing for visual and olfactory contact. The next morning, the divisor was removed to let each pair spawn. Breeding tanks were checked every half hour and eggs were collected and transferred into Petri dishes. They were placed in a E3 and 0.1% methylene blue solution to avoid fungal growth and placed in an incubator set at 28°C.

### Computer Assisted Sperm Analysis (CASA)

To assess sperm motility, we used Computer-Assisted Sperm Analysis (CASA; ISAS; Proiser, R+D, S.L.). We placed 2 µl onto a mixing plate which was activated with 3 µl of tank water at 28°C on a Cytonix 4 Chamber slide (MicroTool B4 Slide, 20 µm depth). A recording taken every ten seconds starting 10 seconds post activation until 60 seconds post activation. We recorded sperm movement using a brightfield microscope (UOP UB203i trinocular microscope; Proiser) at 100× magnification and a black and white video camera (782M monochrome CCD progressive camera; Proiser). The recordings were analysed using ISAS v1 software (Proiser) with the following settings: frame rate: 50 frames/s; frames used: 50; particle area: 5–50 μm2; threshold measurements for VCL: slow, 10–45 μm/s; medium, 45–100 μm/s; rapid, >100 μm/s.

### Embryo/larvae phenotype

We placed individual fertilised eggs onto 12-well plates for monitoring of hatching rate every two hours from 48 hpf, yolk utilisation and growth rate at 24 hpf and between 5 and 8 dpf. A picture of each embryo was taken under a dissection microscope at 24 hpf, 5dpf and 8dpf and the measurements were taken for yolk diameter at 24 hpf, larval length, yolk length and lipid droplet at 5dpf and of larval length at 8dpf. Images were quantified using ImageJ software and scale bar for reference.

### RNA extraction

For transcriptome analysis, we used one to three 24-hpf embryos from each clutch that were manually dechorionated and flash frozen in 1.5 ml Eppendorf tubes. RNA extraction was performed with the RNeasy micro kit (Qiagen) following the manufacturer’s instructions. Briefly, samples were lysed in RLT buffer passing the embryos through a needle and syringe. Lysates were passed through a MinElute spin column and centrifuged for 15s at 13000 g. samples were washed with buffer RW1 and treated in-column with DNase I diluted in buffer RDD for 15 minutes at room temperature (RT). The digestion reaction was stopped with RW1 buffer and centrifuge step. Columns were washed with RPE buffer and 80% ethanol followed by a 5 minute centrifuge step to remove any ethanol residues. Finally, we eluted the RNA in 14 μl of elution buffer and proceeded with quality assessment by TapeStation HS RNA tapes (Agilent) **(Table S1)**.

### Library preparation and RNA-seq

The library preparation was performed with the Lexogen QuantSeq 3’ mRNA-Seq Library Prep Kit FWD for Illumina (Lexogen). RNA was normalised to 13.5 ng/μl and the RNA input for prep = 67.5 ng per sample. The protocol for high quality samples was used. SIRV set 3 were spiked into the samples at a very low level to allow technical evaluation of library prep and sequencing performance. 18 cycles of PCR were performed. Libraries were quantified with Picogreen assay (Fisher Scientific) and sizes were determined with a HS D1000 tape (Agilent) **(Table S2).**

Libraries were pooled to 4nM final concentration and spiked with 1% phiX and processed on a NextSeq flow cell (150 cycles mid output) for sequencing on a NextSeq 500.

### RNA-seq analysis

Following Lexogen guidelines, sequencing adapters, polyA read through and low-quality tails were removed by using the *bbduk* tool of the *bbmap* package (parameters: k=13, ktrim=r, useshortkmers=t, mink=5, qtrim=r, trimq=10, minlength=20) (Bushnell, 2014). RNA-seq reads were then mapped to the zebrafish reference genome (GRCz11.96) using STAR with default parameters (version: 2.6.0c) (10.1093/bioinformatics/bts635). Gene expression levels were estimated by the read counting module embedded within the STAR tool (–quantMode GeneCounts). Differentially expressed genes were identified by using edgeR (10.1093/bioinformatics/btp616) using the TMM method to normalise raw read counts, whereas common, trended and tagwise dispersions were estimated by maximising the negative binomial likelihood (default). Lowly expressed genes were removed (only genes expressed >1 TMM in at least 50% of the samples were retained) and differentially expressed genes were tested performing quasi-likelihood F-tests (glmQLFit and glmQLTest), including the maternal genetic background of each embryo pair in the design of the linear model. Only genes showing FDR < 0.1 were considered as differentially expressed. GO enrichment analysis was performed by using topGO (Alexa & Rahnenfuhrer, 2022) on the GO terms associated to the up- and down-regulated genes separately. In both analyses, GO terms associated to genes expressed >1 TMM in at least 50% of the samples were used as background. The statistical significance of the enrichments was tested with a Fisher’s Exact Test (algorithm=’weight’), then, GO terms associated to less than two significant genes were discarded prior to FDR calculation (Benjamini & Hochberg). Significant threshold was imposed to FDR < 0.1.

The gene co-expression network was built with BioLayout V. 3.4 using expression values for the deregulated genes as input. The Pearson correlation cut-off was set to 0.94 and Markov clustering (MCL) approach was used to determine clusters of co-expression. The protein-protein interaction (PPI) network was built on STRING (V. 11.5) using the ensemble gene IDs of the DEGs and calling clusters of interaction through MCL with inflation parameter set to 2. KEGG pathway and anatomical term enrichments of the PPI network clusters and DEG, respectively, were studied with ShinyGO. The chromosomal ideogram was made with Phenogram (Wolfe et al., 2013).

The intersection between DEGs in intestines of starved adults (Jawahar et al., 2022) and the offspring of starved males (this study) was done using the merge function in R. The odds ratio of the common expressed genes in the two datasets was calculated with using medcalc (https://www.medcalc.org/calc/odds_ratio.php).

Single-cell RNA-seq data from 24 hpf embryos (Lange et al., 2022) were used as reference to deconvolute anatomical information from our DEGs detected upon bulk RNA-seq data analysis. We followed the detailed protocol in Marquez-Galera et al (2022) (Marquez-Galera et al., 2022). Briefly, the anndata object was converted to a h5Seurat file and read into a seurat object. Single-cell data counts were normalised and PCA, and TSNE dimensionality reductions were applied. DEGs from bulk RNA-seq were then read and intersected with the single-cell data and shifts in the cell populations in the dimensionality reductions were used as indicator of populations enriched on these gene sets.

### Statistical analysis

All analyses were conducted using R v. 4.1.2. Linear models were fitted using the lm function and Generalised Linear Mixed-Effects models were applied using the glmer function from lme4 package on phenotypic data following a binomial response (i.e. embryo survival and fertilisation success). Log-transformed sperm VCL was analysed applying the Linear Mixed-Effects Models through the lmer function from lme4. ANOVA type III followed by a post hoc Tukey test were then applied to these models. Graphs for the phenotypic data, KEGG pathways, tissue specificity and transcription factor predicted motifs were produced with the ggplot2 package.

## Results

We exposed adult zebrafish males to one of two treatments: a Control treatment where males were fed an *ad libitum* diet of dry and live (artemia) food three times a day (hereafter referred to as *fed* males) and a short-term Starvation treatment where males were deprived of food for 18 days (hereafter referred to as *starved* males) (**Fig. 1A**). In zebrafish, a full cycle of spermatogenesis takes six days on average (Leal et al., 2009) and the 18 days of starvation allowed for at least two full spermatogenic cycles ensuring the full exposure of germ cells to the treatments. Starved males lost 21.83% of their weight by the end of the experimental period (day 18), an effect that was confirmed in three separate experiments (**Fig. 1B, Fig. S1A**) while fed males showed no significant change in weight. We assessed the effects of starvation on ejaculate parameters as a first indicator of possible effects on the germ cells and reproductive fitness which could lead to intergenerational effects **(Table S3-5)**. We found no significant difference in sperm density between starved and fed males **(Fig. S1B)** but sperm from starved males were overall slower over time (**Fig. S1C-D**). Fertilisation success did not differ between males from the two treatments in *in vitro* fertilisation (IVF) assays (**Fig. 1C**) but it was significantly lower in starved males in natural spawning with wild-type non-experimental females compared to fed males (**Fig. 1D**) together with the total number of eggs laid in their crosses (**Fig. S1E**).

### Paternal starvation effects on offspring survival, growth and reproductive success

We used a split clutch design to disentangle maternal and paternal effects by using the ejaculate of each male to fertilise half of the eggs of two females (**Fig.1A**). When studying offspring survival, we found no significant effect of paternal starvation during the first 24 hours post fertilisation (hpf) or the first 2 months post fertilisation **(Fig. S2A-B)**. However, larvae from starved males were slightly but not significantly smaller at 5 days post fertilisation (dpf) and grew faster (χ^2^=5.1191, DF = 1, p-value = 0.024) compared to their half-siblings sired by fed males ending up significantly larger at 8 dpf showing no other morphological differences. (**Fig. 2A**, an example image of these larvae at 5 and 8 dpf is provided in **Fig. S2C)**.

**Figure 2.**
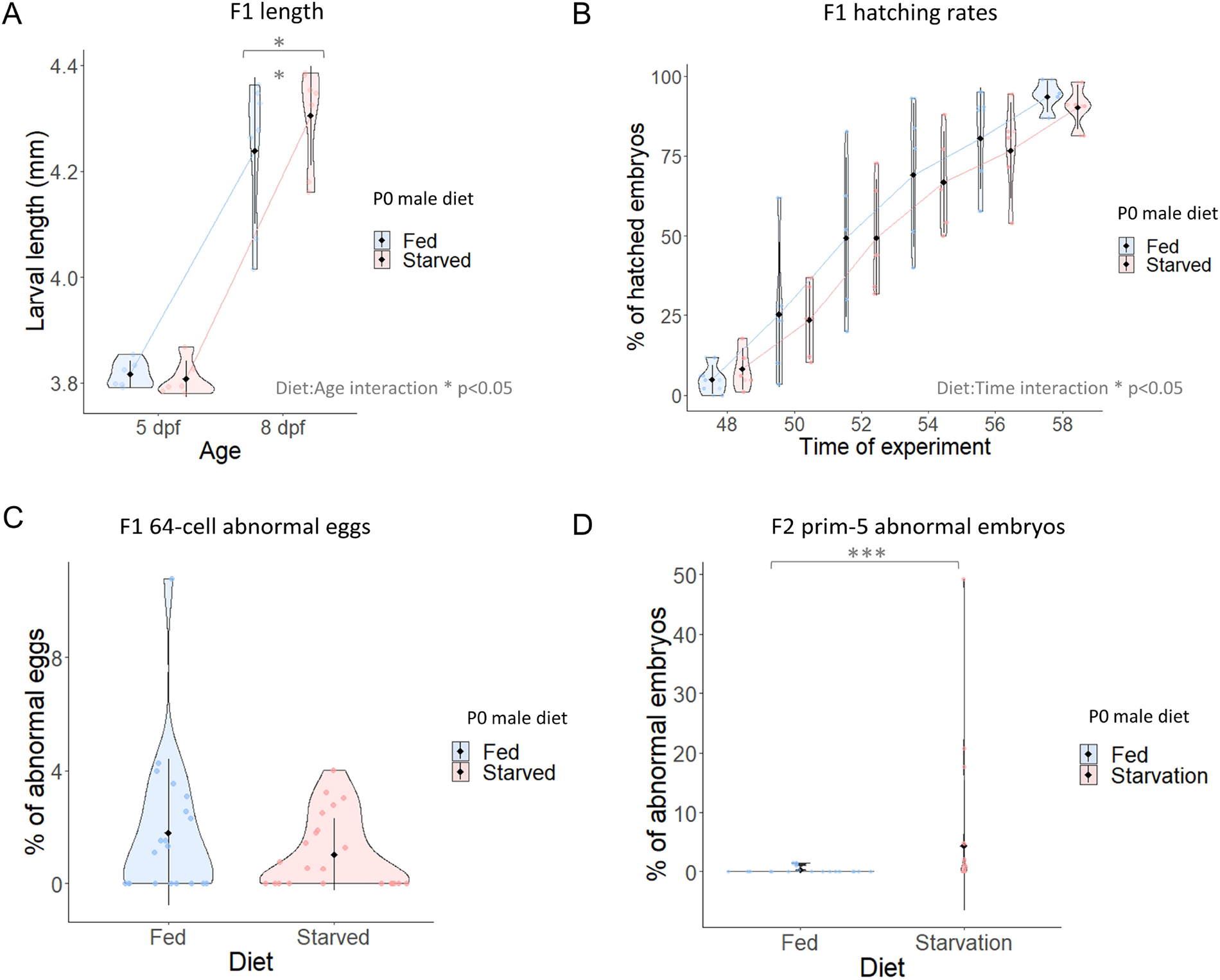
Paternal starvation significantly enhances larval growth but delays hatching and is associated with F2 malformations. **A.** Larvae length: Larvae growth was measured at 5 and 8 dpf in larvae from several clutches. Larvae from starved fathers were significantly longer than their control siblings. Individual data points in the violin plot represent the average of larvae length per IVF (5 dpf: 175 larvae from fed fathers and 180 from starved fathers within 6 independent IVF replicates. 8 dpf: 287 larvae from fed fathers and 278 from starved fathers within 7 independent IVF replicates). **B.** Hatching rates: The percentage of hatched embryos was analysed every 2 hours between 48 and 58 hpf. C. F1 abnormal eggs as determined at 64-cell stage (2 hpf). Eggs from males and females from fed and starved fathers were collected and analysed upon natural mating. No statistical differences were observed in the egg quality between groups or between groups and either sex. **D.** F2 abnormal embryos at prim-5 stage (24 hpf). Increased number of abnormal embryos was observed in the group laid by F1 fish coming from starved fathers. No statistical differences were found when sex was considered as interaction factor.*** p < 0.001; ** p <0.01.

We further assessed hatching rate which is known to be affected by a range of variables during embryo development (Pype et al., 2015; Scopel et al., 2021; Zajitschek et al., 2014). We checked for hatched larvae every two hours between 48 and 58 hpf and found that the total hatching rate was slower in the clutches sired by starved males compared to the clutches sired by fed males (χ^2^=4.6390, DF = 1, p-value= 0.031) (**Fig. 2B**).

During their first days of life, zebrafish embryos rely entirely on the nutrient supply from the yolk which is gradually absorbed until the embryo becomes a free feeding larva at 5 dpf (Anderson et al., 2011; Huang & Linsen, 2015). Therefore, as an indicator of nutrient consumption and metabolic activity, we measured the size of the yolk (length, diameter, and area) at 24 hpf and at 5 dpf. We found no significant effect of paternal starvation on yolk size at either time point **(Fig. S2D-F)**. The difference in growth rate between offspring of starved and fed males therefore appears not to be explained by changes in mobilisation of maternal provisioning during these early stages as measured by yolk size and may reflect metabolic differences in the growing cells of the embryos and larvae.

The sex ratio in the offspring upon reaching sexual maturity did not differ between starved and fed males **(Fig. S2G).** When setting up male and female F1 offspring from starved and fed males with wild-type non-experimental fish for natural spawning we found no overall difference in fertilisation success, abnormal eggs or offspring survival during the first 24hpf **(Fig. S2H, Fig. 2C, Fig.S2I)**. However, we found a significantly higher percentage of abnormally developing F2 embryos at 24hpf produced by the F1 offspring of starved males compared to the clutches produced by the F1 offspring of fed males (**Fig. 2D**).

### Impact of paternal diet on the F1 transcriptome

To trace the potential molecular changes underlying the observed phenotypic effects of paternal diet on early embryo development and adult reproductive fitness in the offspring, we explored transcriptome of prim-5 embryos (24 hpf), a stage of development characterised by fast growth and dynamic expansion of transcriptome repertoire (White et al., 2017), which represents the most conserved transcriptomic stage among anamniotes fitting with the developmental hourglass model (Marlétaz et al., 2018). Prim-5 embryos from starved and fed males were generated in split-clutch IVF assays as described above.

Transcriptomic data of embryos shows great variation beyond the feeding differences (**Fig. S3A)**. Yet, despite the variation in these embryo transcriptomes our analyses revealed distinguishing changes both in the down and upregulated sets due to paternal diet in the transcriptome profile of the F1. Indeed, differential gene expression analysis of 3’ end transcript identified 145 differentially expressed genes (DEGs) between embryos from starved and fed males (**Fig. 3A-C**). The majority (115 genes) of the DEGs were downregulated in offspring from starved males while a smaller subset (30 genes) was upregulated **(Full details provided in TableS6)**.

**Figure 3.**
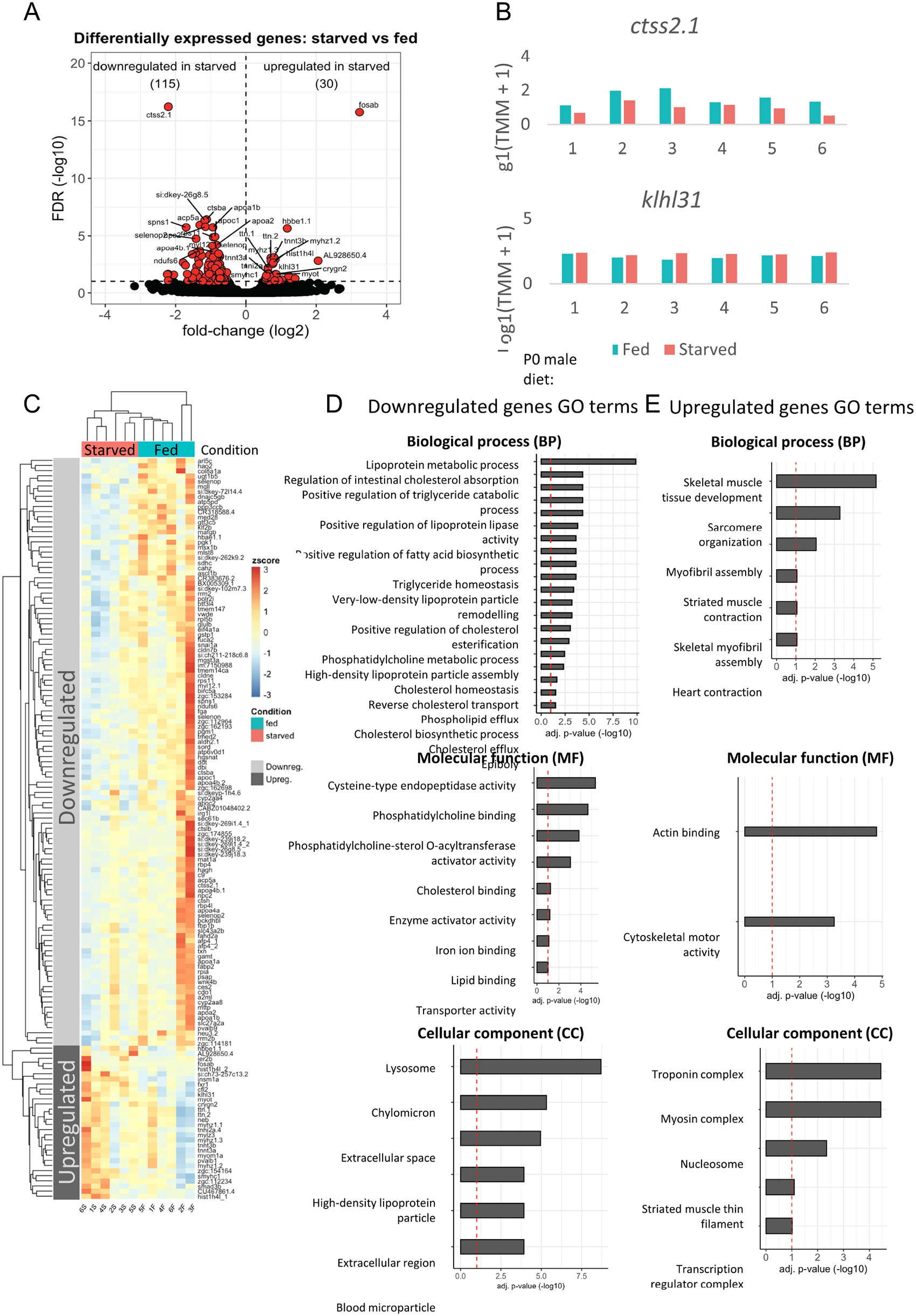
Paternal starvation leads to changes in embryo transcriptome in prim-5 embryos. **A.** Differentially expressed genes in offspring of starved and fed fathers: Volcano plot depicting the differentially expressed genes between embryos from starved and fed fathers. We found 130 genes downregulated and 15 genes upregulated (only genes expressed >HMM in 50% of the samples, FDR<0.1). A total of 6 samples for each group were included in the analysis. **B.** Example of expression levels in downregulated (top: *ctss2.1)* and upregulated (bottom: *klh/31)* genes in offspring of starved fathers. Examples are given for all the embryos included in the transcriptomic analysis. **C.** Differential expression within experimental groups and deregulated genes: Heatmap showing the variation in downregulated genes (top) and upregulated (bottom) between embryos from starved and fed fathers. **D.** GO terms for upregulated genes in embryos from starved fathers. **E.** GO terms for downregulated genes in embryos from starved fathers.

To control for transcriptomic bias given by the samples used as reference in the analysis, we compared the transcriptome of embryos from fed males to the transcriptome of an independent set of 24 hpf wild-type embryos provided by a different fish facility but processed following the same protocol as the embryos from our starvation experiments and included in the same sequencing run (**Fig. S3B**). This analysis showed a strong correlation between the transcriptomes from the embryos from fed males in our experiments and the independent wild-type embryos confirming the validity of our findings of DEGs and the robustness of our results.

Next, we sought to understand the biological relevance of the observed gene expression changes. A GO term analysis of the 115 downregulated genes revealed significant enrichment for genes associated with lipid metabolism and lysosome (GO terms such as “lipoprotein metabolic process”, “regulation of triglyceride catabolic process”, “positive regulation of fatty acid biosynthetic process”, “cholesterol homeostasis”, “cholesterol binding”, “lysosome”) (**Fig. 3D**). In contrast, the GO term analysis of the 30 upregulated genes showed an enrichment for genes related to muscle development and contraction (GO terms such as “skeletal muscle tissue development”, “sarcomere organization”, “myofibril assembly”) (**Fig. 3E**). Then we asked whether the genes showing similar trends of regulation are potentially co-regulated for example by chromosomal topological constraints. Gene location analysis indicated that DEGs were distributed across the entire genome, but we found small clusters of genes located on chromosomes 5 (*myhz* and *c9*), 12 (*ctsl*) and 16 (*apoa* and *apoc*) (FDR<0.01) **(Fig. S4)**. These genes are related to muscle development, cellular response to estrogen stimulus, proteolysis, and lipid transport; respectively. The deregulation of these clusters suggests a potential co-regulation of the expression of these genes. We explored this further by building a gene co-expression network to which we applied a Markov cluster algorithm (MCL clustering). This showed that several of the genes clustering together on a chromosome were also found within the same co-expression **network clusters** (Fig. S5A-C Clusters 1, 4 and 5. A full list of the genes included in each cluster is provided in TableS7).

To further understand the relationship between the DEGs and their gene products, we created a protein-protein interaction (PPI) network and determined clusters of interaction by applying the MCL clustering approach. Based on known and predicted interactions, this analysis revealed nine clusters of interacting proteins for the downregulated genes (Clusters 1 to 5 showed enriched GO terms for lipoprotein metabolic process, proteolysis, monosaccharide metabolic process, ribosomal large subunit assembly and deoxyribonucleotide metabolic process, respectively) and two interaction clusters for the upregulated genes (cluster 1 enriched for GO terms associated to muscle contraction and cardiac muscle contraction, among others) **(Fig. S6A-B)**. The analysis of the KEGG pathways enriched within these clusters revealed terms such as PPAR signalling and autophagy (**Fig. 4A**) whose activity is related to the nutritional status of the cell including lipid metabolism. We found no KEGG pathways significantly enriched for the interacting genes in clusters 1 or 2 of the upregulated genes PPI network.

**Figure 4.**
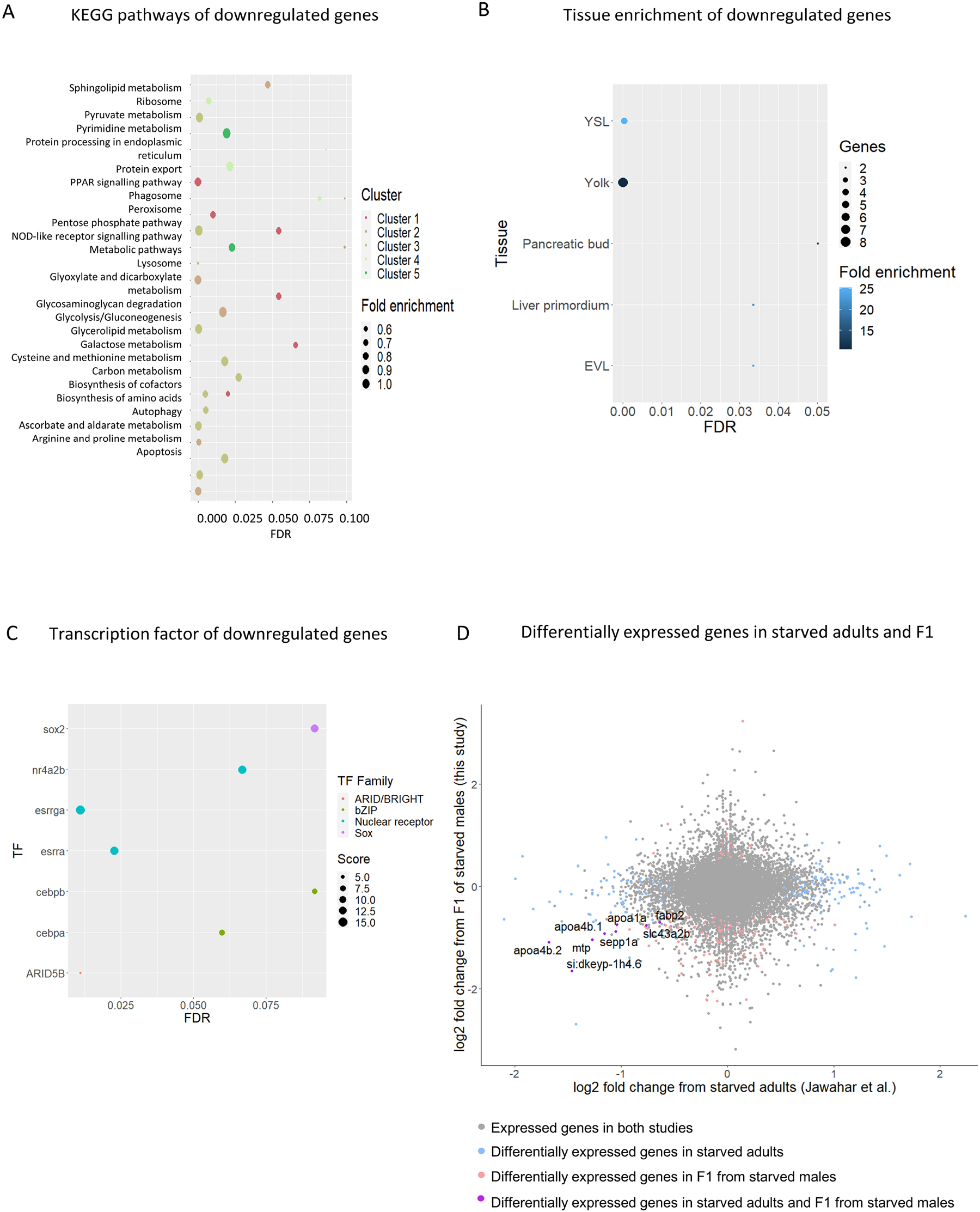
The downregulated genes in the offspring of starved males are associated to metabolic processes, show preference for yolk and YSL expression and correlate with altered genes in adults exposed to starvation. **A.** KEGG pathways enriched in downregulated genes. **B.** Tissue enrichment of downregulated genes. **C.** Transcription factor enrichment in the subset of downregulated genes. **D.** Intersection analysis between the DEGs in the datasets from Jawahar et al. (2022) and this study (Odds ratio= 4.9617, p<0.0001).

When interrogating the tissue localisation of the pool of downregulated genes based on the anatomy information in the generif database compared to a background list of the total genes expressed at 24 hpf, we found significant enrichment in the yolk syncytial layer (YSL), yolk, the enveloping layer (EVL), pancreatic bud and liver primordium (**Fig. 4B**). We sought to further explore the tissue specificity of expression of DEGs at 24 hpf. To this end, we used existing single-cell RNA-seq dataset produced by the Zebrahub partners (Lange et al., 2022) to which we applied a dimensionality reduction approach (Marquez-Galera et al., 2022) to differentiate between cell-type-specific signals in bulk RNA-seq DEGs according to the single-cell RNA-seq data at 24 hpf. This approach allows us to increase the accuracy of the spatial distribution of the DEGs as well as the specificity to developmental stages **(Fig. S7).** This analysis revealed an enrichment of the downregulated genes in cells associated with the hematopoietic system, blood island, hatching gland and periderm (tissue derived from the EVL) (**Fig. S7 mid panel**). Note that the clusters defined in the single cell data lack information for pancreatic bud and liver primordium. These are small cell populations which might be more difficult to pool in this type of approach. The absence of yolk and YSL information is given by the nature of the sample preparation for single cell RNA-seq where embryos are dissociated, and samples are washed and spun leading to loss of the lipid-rich yolk in the supernatants; or yolks are directly removed with deyolking buffer (Farnsworth et al., 2020; Jiang et al., 2021).

Because of the lack of reference transcriptomic information for the yolk in the single cell RNA-seq, we then sought to further confirm the findings provided by generif by searching the *in-situ* hybridisation data in the zebrafish information network database (zfin) and associated available literature. This manual curation of the data allowed us to confirm the results reported by the generif database for all the YSL expressed reported genes (i.e. *fabp2, mttp, apoa4b.1, cldne, gamt, apoc2, apoc1, rbp4)* and added information on YSL expression on 29 additional genes **(Table S6).** The yolk and YSL are essential extra-embryonic structures involved in the metabolic activity of the early embryo. The downregulation of these genes in the yolk and YSL in embryos from starved fathers points to the impact of paternal nutrition on these tissues with possible impact on the metabolic activities of the offspring. The generif approach revealed no significant enriched tissues for the upregulated genes. However, the deconvolution of single cell data with the upregulated genes showed that those genes are expressed in cardiac muscle cells, myotome, notochord, and muscle pioneers at 24 hpf (**Fig. S7 bottom panel**). These results were further validated by the data available in zfin (Bradford et al., 2022).

Furthermore, when we explored the transcription factors (TFs) predicted to bind to the downregulated DEGs, we found enrichment for *cebpa* and *cebpb*, transcription factors among others. These factors, members of the bZIP family, are related to the digestive system development (Yuan et al., 2015)(Yuan et al., 2015) and have enriched expression in the YSL, liver and gut (**Fig. 4C**) again pointing at the misregulation of genes involved in nutrient utilisation.

Finally, we queried the relationship between transcript abundance profiles upon direct exposure to starvation and offspring response. To this end, we compared the transcriptome data from the embryos from starved males in our study with existing transcriptome data from intestines of starved adult zebrafish (Jawahar et al., 2022) (**Fig. 4D**). This dataset offers a good opportunity to focus on the impact of the dietary stressor which in our unexposed embryos seems to target a nutritionally relevant tissue in the early embryo as observed by the pool of deregulated YSL-associated genes. This transcriptome intersection analysis revealed a significant enrichment for a subset of genes that were downregulated in both studies (Odds ratio = 4.9617, p<0.0001). These genes (*apoa1a*, *apoa4b.1*, *apoa4b.2*, *fabp2*, *mtp*, *sepp1a*, *slc43a2b*, *si:dkeyp-1h4.6*) belong to cluster 1 in our PPI network and are mostly involved in PPAR signalling. This finding indicates that the impact of starvation can affect the same metabolism pathway genes similarly in starved adults and in unexposed offspring.

## Discussion

Parental condition is known to impact the fitness of the following generations (Bautista et al., 2023; de Assis et al., 2012; Xu et al., 2023) and maternal effects are well characterised (Jung et al., 2022; Ondičová et al., 2022). In contrast, paternal effects have only recently been recognised as biologically relevant. One reason for this is that separating the paternal effects from maternal confounding factors is often difficult (Lacal & Ventura, 2018). In this study, we used a split-clutch approach that controls for maternal contribution through the comparison of maternal half-siblings in a zebrafish model. Our results provide clear evidence that paternal starvation not only affects male fertility but also directly affects offspring performance early and later in life by impacting the development of the F2 generation.

We found that reduced swimming velocity was exhibited by sperm from starved males, but interestingly, this only affected fertilisation success during natural mating. Nevertheless, the finding of reduced velocity in sperm suggests that starvation has an impact on spermatogenesis and affects the cellular processes, potentially by affecting metabolic pathways that also determine sperm velocity. Interestingly, fertility traits showed great variability among starved males suggesting that the overall condition of the males may interfere with the impacts of starvation, and that males that are generally in better condition may be more resistant. Genetic variation among males is the most likely explanation for the observed variation in our experiment, as a such variation was reported from recombinant inbred mice exhibiting marked differences in their response to dietary restriction (Liao et al., 2013). The potentially condition-dependent response to dietary restriction needs further investigation.

The changes triggered by paternal starvation leading to reduced fertility and sperm fitness can be linked to non-genetic alterations transmitted to the offspring. When looking at the impact of this treatment on early life performance in the resulting offspring, we found that while survival during early development and juvenile life stages were not significantly affected, hatching rate was slower and growth rate faster in larvae sired by starved males compared to their half-siblings sired by fed males. Despite it not being largely different, the finding of faster growth was highly reproducible and is similar to those alterations observed upon maternal starvation in the offspring of *C. elegans* (Hibshman et al., 2016) or in mice with overgrown pups produced following reduced paternal protein diet and caloric restriction (Morgan et al., 2021). Nevertheless, our results contrast with the reduced growth in larvae as a result of maternal starvation in zebrafish (Fan et al., 2019).The inconsistency on outcomes upon parental caloric restriction suggests that the impact of parental diet is highly influenced by the conditions of the model, including species, age, sex and duration of exposure to the stressor. For example, fish undergo long periods of starvation that can exceed weeks and are more resistant to this condition (Drew et al., 2008; Furné et al., 2012; Furne & Sanz, 2023; Wang et al., 2006). Hence, what we consider as short-term starvation in our study could represent an intense exposure in other species.

The faster growth rate during early larval stages suggests more efficient utilisation of the available resources towards muscle formation and may translate into potentially negative effects later in life in reproductive fitness, as indicated by the overall higher rate of abnormally developing embryos produced by the offspring of starved fathers. The integrity of germ cells is costly, and its maintenance is often at the expense of the soma (Chen et al., 2020; Ivimey-Cook et al., 2022). The decay in reproduction fitness with enhanced growth observed in offspring from starved fathers could be either a cause or a consequence of a trade-off between soma and germline as influenced by the paternal environment and manifested early in development. These findings are interesting as they suggest that while short-term starvation in males may be beneficial for individual lifespan and health, they may have detrimental effects both on individual reproductive fitness and adult condition and reproduction in the resulting offspring.

In our study, yolk size did not differ between embryos of starved and fed males meaning that the observed difference in growth rate was not directly linked with resorption of yolk which otherwise is indicative of metabolic turnover (Anderson et al., 2011; Huang & Linsen, 2015). However, the finding of a total of 145 DEGs several of which robustly associated with major functions in body growth and fatty acid metabolism further supports the idea that the paternal condition directly influences gene expression in the offspring. Several metabolism-related pathways were enriched in the protein-protein interaction clusters formed by the downregulated genes in this analysis and included PPAR signalling, autophagy and glycolysis/gluconeogenesis. These pathways are often found deregulated in conditions of direct exposure to starvation. For example, autophagy is triggered in conditions of nutrient deficiency upon mTOR pathway inhibition (Mizushima et al., 2004; Wong et al., 2015) while *pparα* is overexpressed under fasting conditions (Kersten et al., 1999).

The mis-regulation of starvation-associated pathways and similitude of the pool of genes affected suggest a remarkable similarity of response in the offspring to that in the parent. This idea is further supported by the significant correlation with data from intestine of starved zebrafish adults (both males and females) (Jawahar et al., 2022) and the transcriptome of our unexposed embryos with a high proportion of misregulated genes being expressed in a nutrition-related tissue like the yolk. The deregulation of similar genes and pathways between parents and offspring has been described in other studies but the response is often opposite to that observed in the parents (Carone et al., 2010; Fan et al., 2019). In our study, we report both: On one hand, genes related to *ppar* signalling mimic the direct response to a similar starvation paradigm. On the other hand, autophagy and other metabolic-related genes show the opposite response to the expected upon starvation. This contrasting behaviour of gene expression could be behind the desensitisation to the stressor exhibited by the offspring of parents exposed to similar conditions, as shown by increased resistance to starvation in the offspring of C. elegans upon maternal starvation (Hibshman et al., 2016). Further experiments are required to investigate whether this is the case in our model but this comparable response to paternal diet between directly exposed adults and non-free feeding and unexposed offspring suggests a functional link to the paternal nutritional status and a directed transmission of response. This directed transmission of paternal information into the offspring could be instructive for the embryo on responses to nutrient availability in the environment despite of the actual nutritional status, which in our experiments was controlled by the mother through the egg content.

Interestingly, we observed that a number of DEGs were located in proximity forming clusters in different chromosomes (5, 12 and 16). These genes respond similarly to paternal starvation. These two facts (clustering and unified response) could point to a co-regulation phenomenon behind the expression of these genes which seem to go hand-by-hand. This co-expression of neighbouring genes has been widely studied and confirmed across several models (Fukuoka et al., 2004; Purmann et al., 2007; Williams & Bowles, 2004) and could point to a conserved response to caloric imbalances.

We also applied different approaches to study the tissue specificity of the DEGs. The upregulated genes showed clear enrichment for GO terms and pathways related to muscle formation and contraction. These genes are mostly expressed in tissues related to these activities (i.e muscle pioneers, notochord and cardiac muscle cell). We speculate that the significant enhanced growth observed at 8 dpf could be a product of the transcriptional deregulation initiated at earlier stages and detected by our transcriptomic analyses at 24 hpf.

The observed downregulated genes are mostly expressed in the yolk and YSL, periderm, hematopoietic system, blood island and hatching gland at 24 hpf. The yolk is an extraembryonic tissue loaded with maternally supplied factors, that operates as the metabolic hub of the early zebrafish embryo and provides nutritional support during the first days of life (Anderson et al., 2011; Huang & Linsen, 2015). Many of these yolk-associated genes are also expressed in the yolk sac of other vertebrates including humans, mouse and chicken (Cindrova-Davies et al., 2017) and their physiological roles in the yolk sac are highly conserved during the early stages of life, meaning that the metabolic response to paternal starvation could also be conserved in higher vertebrates. The altered gene expression caused by paternal starvation in this tissue could translate into metabolic defects which ultimately affect larval growth and fitness. However, metabolomics studies would be required to test this hypothesis.

The potential mechanisms that could lead to altered gene expression in extraembryonic and embryonic tissues are various and complex and include changes in DNA methylation, histone modifications and small RNAs. For example, high fat diet alters H3K4me3 deposition in sperm in genes associated to placental formation (Pepin et al., 2022). On the other hand, tRNAs and miRNAs carried in the ejaculate can change upon protein restriction leading to transcriptomic changes in the offspring (Sharma et al., 2016). However, in externally fertilizing fish, the impact of the ejaculate might be reduced by the dilution of its contents in water (Fitzpatrick, 2020).

As the majority of the mis-regulated genes in our study appear downregulated, it is possible that the mechanisms that mediate paternal starvation effects on offspring transcriptome at the studied stage include a combination of these since, for example, the direct deposition of H3K4me3 in sperm in the metabolic pool of genes would not explain the decreased expression observed here.

### General conclusions

Overall, our study confirms the importance of paternal condition for offspring fitness from early development into late life and across at least two generations. The non-genetic alterations caused by paternal diet in the offspring can have adaptive or maladaptive consequences which are in turned determined by shifts in the environment, among other factors (Stajic & Jansen, 2021). The similarity of the gene expression patterns in offspring of starved males and starved adult males (Jawahar et al., 2022) suggests that the observed response could be potentially adaptive. Whether this is truly the case or whether the inheritance of epigenetic factors from fathers is more of a side effect still needs to be more carefully tested.

## Ethics permit

All animal work was performed under the Project Licences # P0C37E901 and P51AB7F76, in accordance with the UK Home Office regulations and UK Animals (Scientific Procedures) Act 1986, and under Licence C200315/16 of the Swedish Board of Agriculture.

## Acknowledgments

This work was funded by a Human Frontier Science Program research grant (HFSP R0025/2015). AJG is currently supported by the PrecisionTox consortium. We thank Genomics Birmingham for sequencing; the BMSU facility at the University of Birmingham, SciLifeLab zebrafish platform at Uppsala University and the Controlled Ecology Facility (CEF) at the University of East Anglia (UEA, UK) for the animal care; the Zebrahub partners for allowing us to use their single-cell data for our transcriptomic comparisons; Bradley Cairns for his intellectual contribution to the original project and Rui Monteiro for his feedback on the manuscript.

