## Supplementary Figures 1-7 and Supplementary Tables 1-5 for "Paternal starvation affects metabolic gene expression during zebrafish offspring development and life-long fitness"

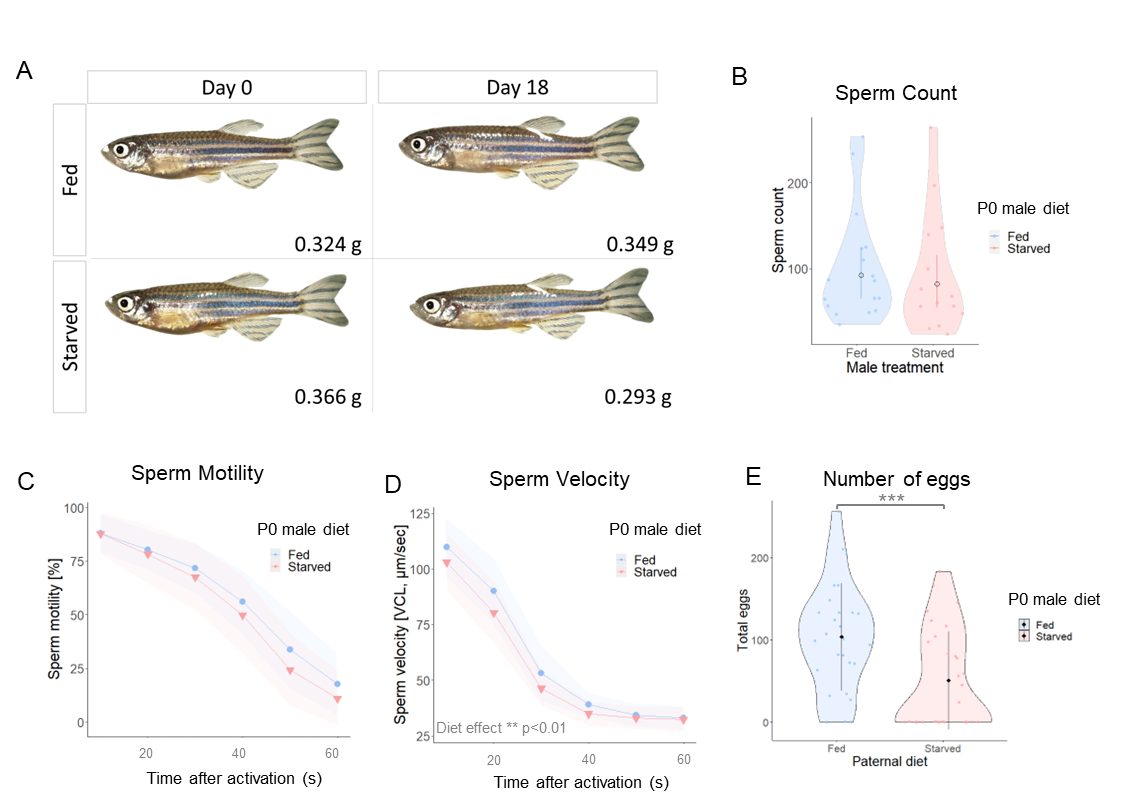

**Supplementary figure 1. Effects of male starvation in body weight and sperm fitness. A.** Example image of fish used in this study. Starved group fish showed consistent reduction of abdominal mass and weight. **B.** Sperm count in fed and starved males. Bars represent the mean and 95% confidence intervals. **C.** Percentage of sperm motility in fed and starved males across 6 time points after sperm activation. **D.** Sperm velocity in fed and starved males at 6 time points after sperm activation. **E.** Number of eggs produced by non-experimental females during natural mating with fed or starved males. Individual data points are represented as dots within the violin plot in panels B and E. Shadowing in panels C and D represents 95% confidence intervals. ** p <0.01. *** p <0.001

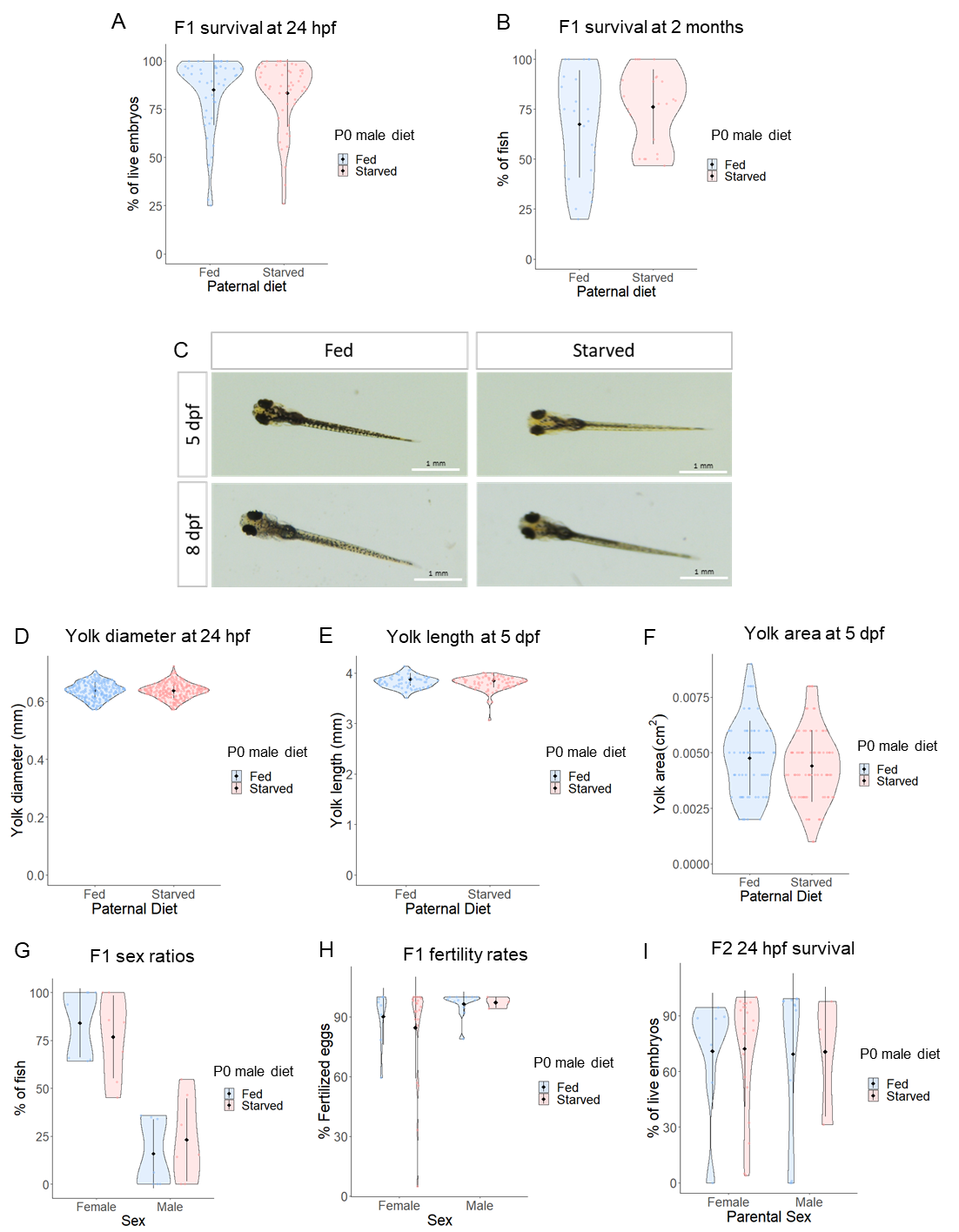

**Supplementary figure 2. F1 phenotypes, fertility assessment and yolk size analysis. A.** F1 survival at 24 hpf: The analysis of several embryo clutches revealed no statistically significant changes in survival between embryos from fed and starved fathers. Individual data points within the violin plot represent the average egg survival rate per male and across 7 independent IVFs (Fed males: 1552 live embryos out of 1777 total embryos. Starved males: 1731 live embryos out of 2038 total embryos). **B.** F1 survival at 2 months post fertilisation. No significant differences were found at this stage. Individual data points within the violin plots represent average offspring survival (321 animals from 7 different IVFs were grown from P generation for each treatment. Survival at 2 months from fed fathers: 235. Survival from starved fathers: 243. **C.** Example images of F1 larvae at 5 and 8 dpf. No evident developmental malformations were observed in the larvae from fathers in the starved group. Black bars represent the mean ± SD. **D.** Yolk diameter of F1 from fed and starved groups at 24 hpf. No differences were observed in yolk size at this stage. **E.** Yolk length in F1 from fed and starved groups at 5 dpf **F.** Yolk area of larvae from fed and starved males at 5 dpf. **G.** F1 sex ratios: We analysed the sex ratios in the offspring from starved and fed fathers. No significant changes were observed. **H.** F1 fertility rates: The study of the fertility rates (Fertilised eggs at 2 hpf) from the offspring of starved and fed fathers bred to wildtype males or females by natural mating revealed no significant changes. Coloured data points represent the average fertilisation rate in eggs from F1 breeding (Fertilised eggs over total laid eggs of Females from fed fathers: 872/944; Females from starved: 1993/2327; Males from fed: 1370/1398; Males from starved: 524/535). **I.** F2 survival at 24 hpf: The analysis of several embryo clutches revealed no statistically significant changes on survival between embryos produced by the F1 (either females or males) of fed and starved fathers. Individual data points within the violin plot represent the average egg survival rate per individual produced by natural spawning.

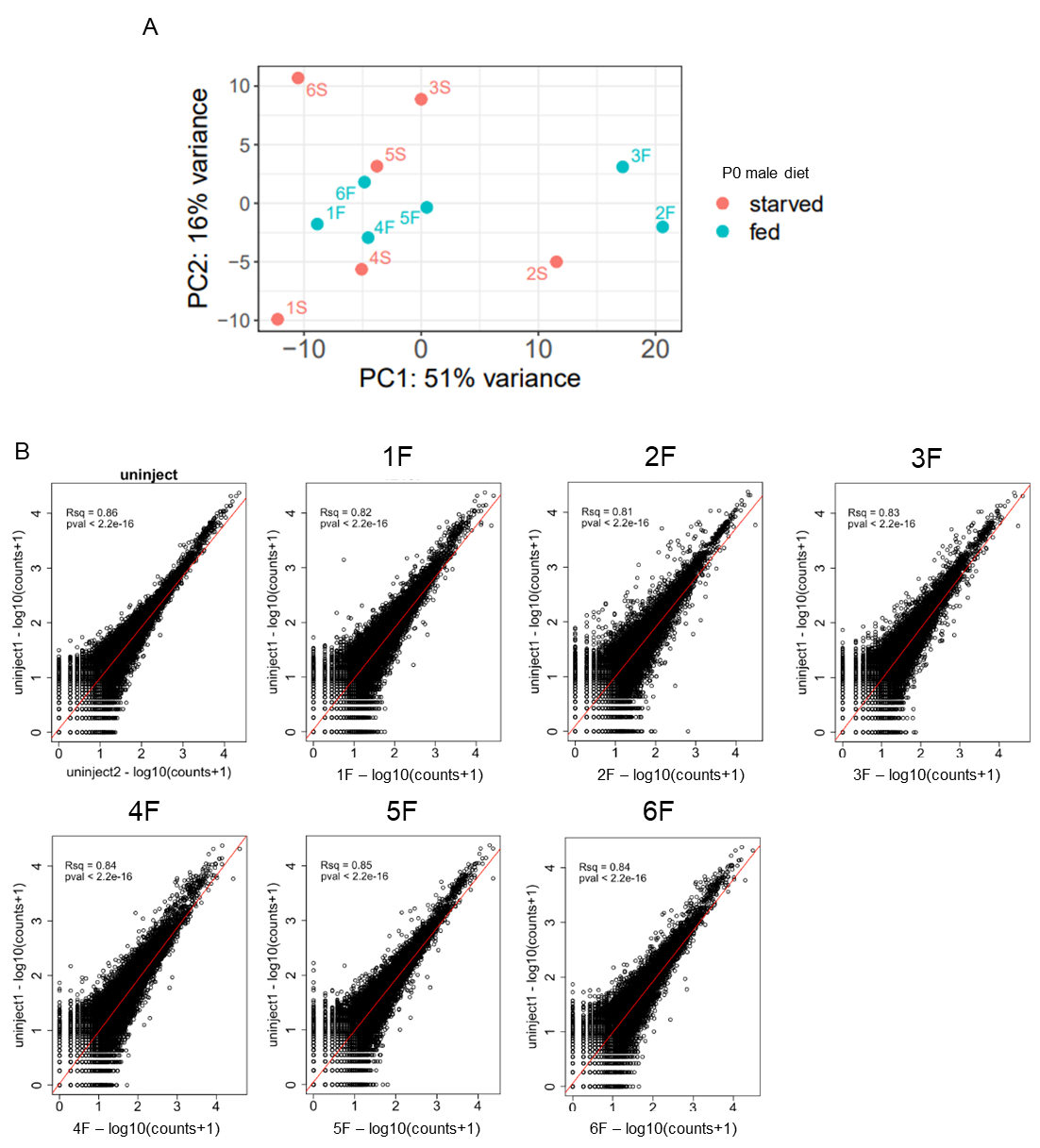

**Supplementary figure 3. RNA-seq controls. A.** PCA plot of gene expression in prim-5 (24 hpf) embryos from fed and starved fathers (N=6, groups contained 1-3 embryos each). **B.** Scatter plots of the correlation between gene expression of AB wt males from a different family, not included in the project and control males used in the starvation experiment.

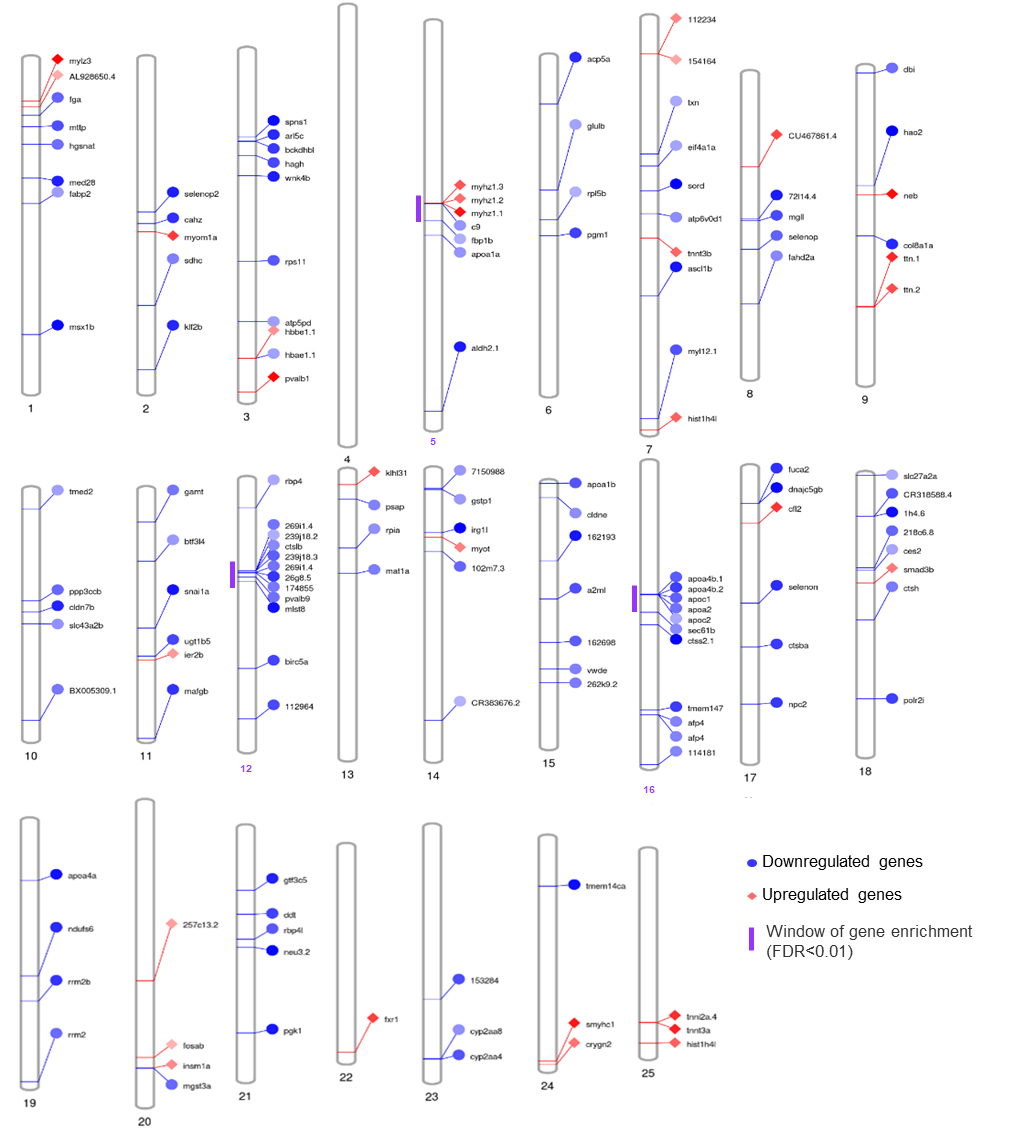

**Supplementary figure 4. Paternal starvation leads to changes in prim-5 embryo transcriptome in genes located in chromosomal proximity.** Ideogram showing the chromosomal localization of the DE genes. We found 3 sets of genes located in clusters in chromosomes 5, 12 and 16. (Purple bars, FDR<0.01)  Blue: downregulated genes. Red: Upregulated genes.

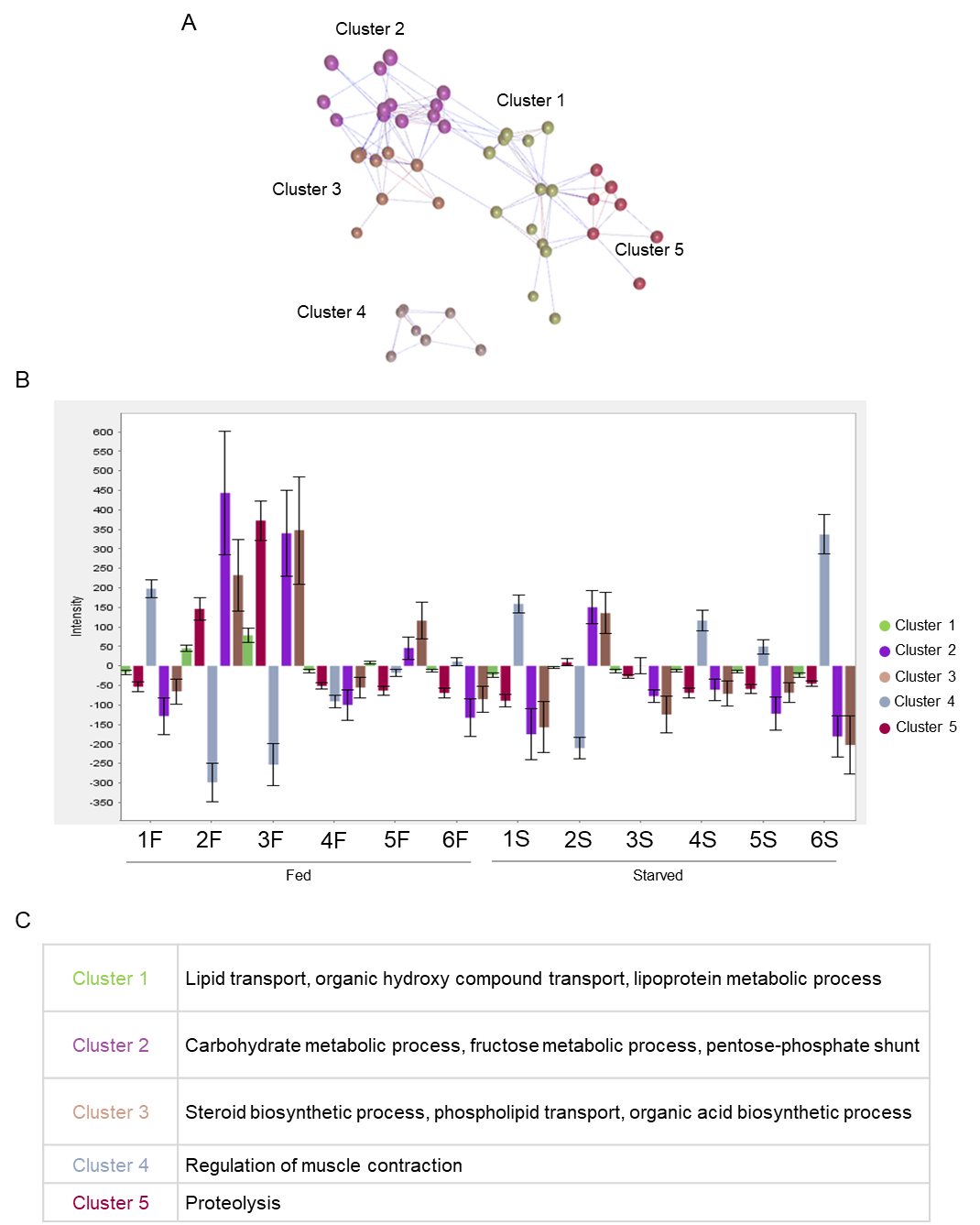

**Supplementary figure 5. Gene co-expression network. A.** Gene co-expression network obtained from gene expression data from the differentially expressed genes in embryos from the starved fathers compared to embryos from fed fathers. Five significant gene clusters were observed. **B.** Cluster expression dynamics across the RNA samples from embryos of fed and starved fathers. **C.** GO terms corresponding to each of the cluster in the gene co-expression network from panel C.

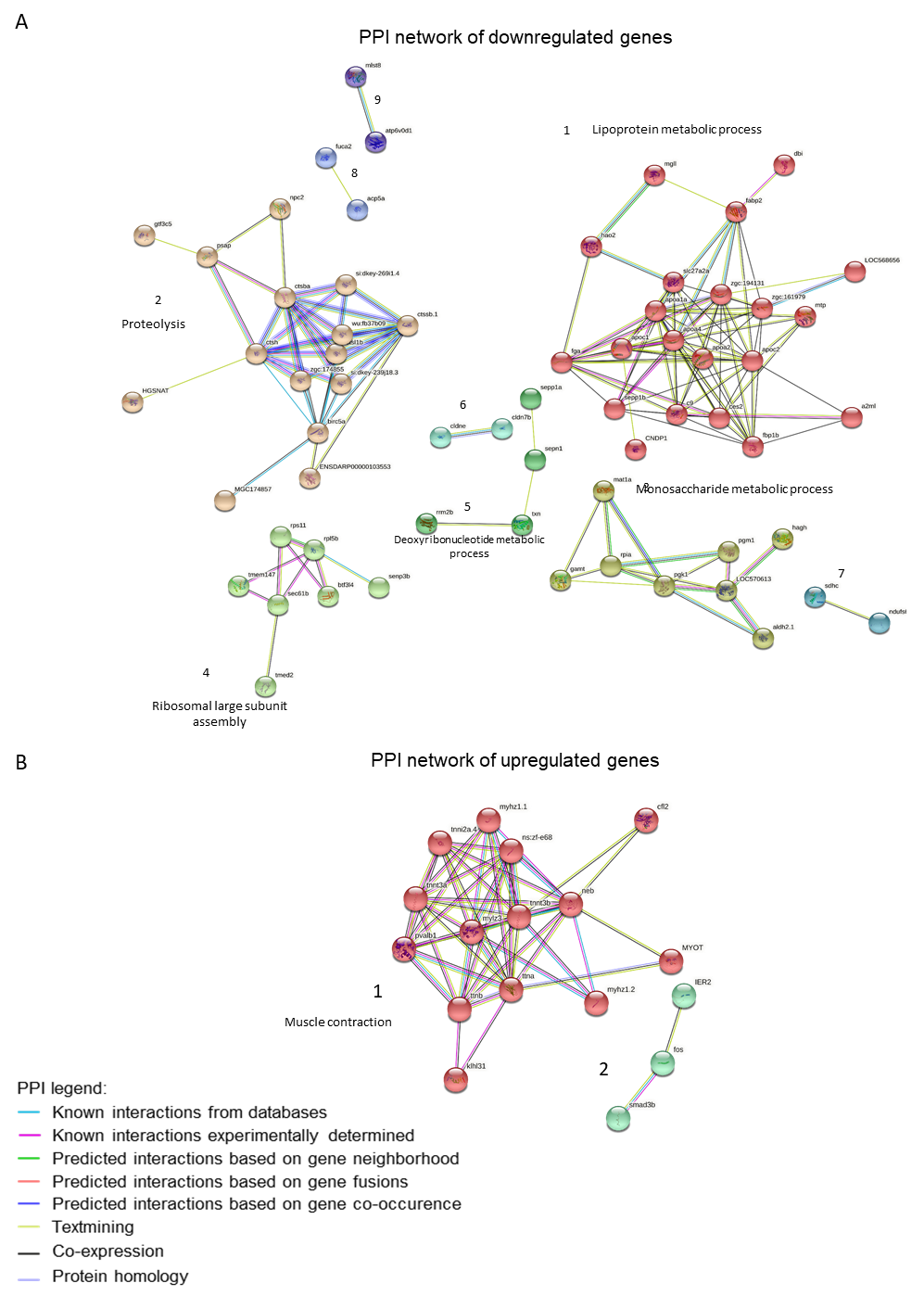

**Supplementary figure 6. A.** Protein-protein interaction network with clustering of the downregulated genes in embryos from starved fathers. **B.** Protein-protein interaction network with clustering of the upregulated genes in embryos from starved fathers.

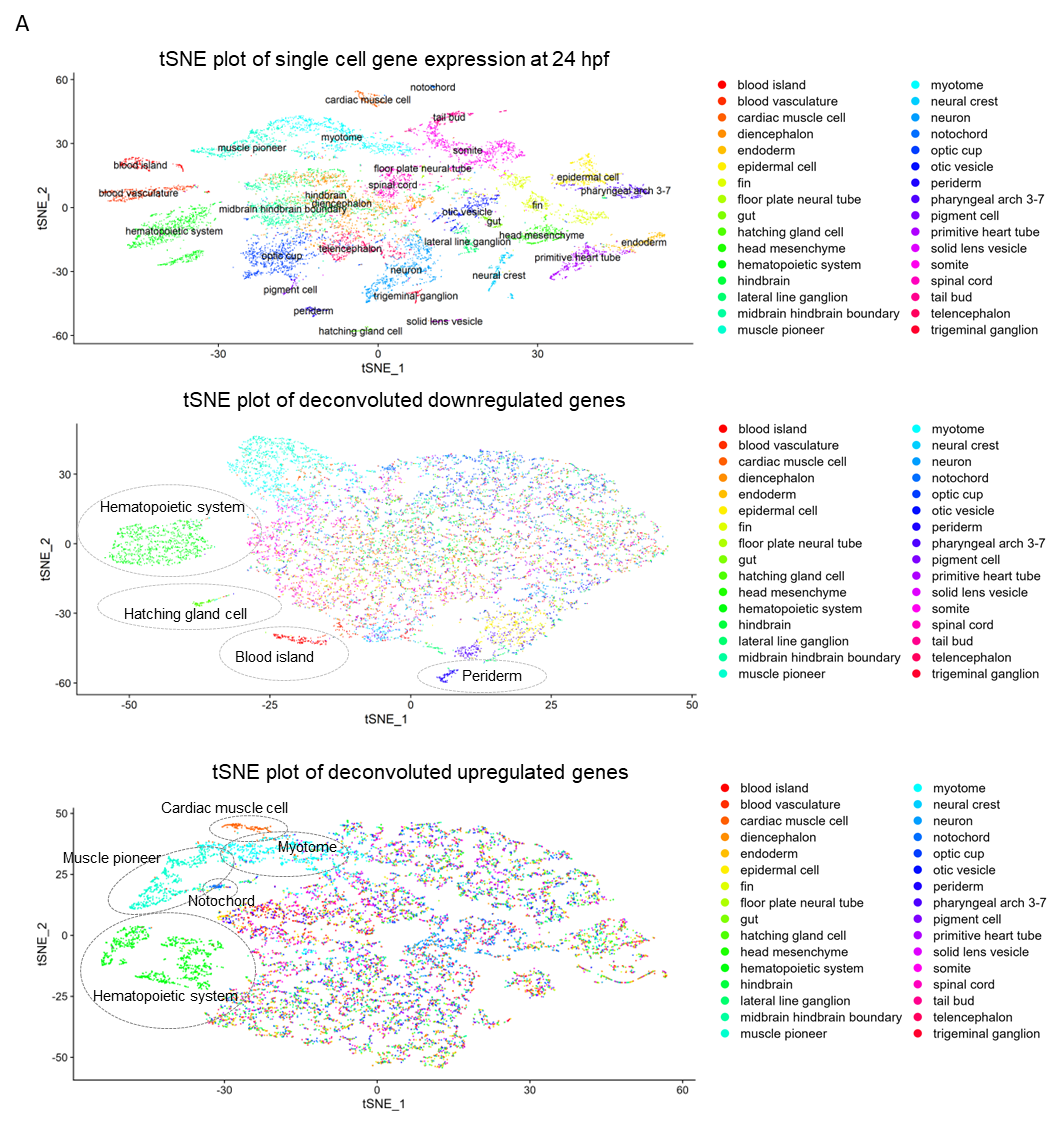

**Supplementary figure 7. Study of the tissue localization of the DEGs upon paternal starvation. A.** Deconvolution of cell-type-specific signal in whole embryo RNA-seq DEGs. Top: tSNE plot of cells types in prim-5 embryos. Middle: tSNE plot of deconvoluted signal from downregulated genes in embryos from starved males with enrichment for hematopoietic system, hatching gland cell, blood island and periderm compatible cells. Bottom: tSNE plot of deconvoluted data from upregulated genes in embryos from starved group males with enrichment for muscle pioneer, notochord, cardiac muscle cell, myotome and hematopoietic system compatible cells.

Supplementary Table 1. RNA integrity number (RIN) of the samples used in the transcriptome study of the F1 generation.

| Sample ID | RIN value |
| --- | --- |
| 1F | 8.3 |
| 1S | 8.2 |
| 2F | 9.3 |
| 2S | 9.3 |
| 3F | 9.3 |
| 3S | 9.7 |
| 4F | 7.2 |
| 4S | 9.1 |
| 5F | 8.9 |
| 5S | 8.9 |
| 6F | 4.8 |
| 6S | 9.8 |

Supplementary table 2. Concentration, library size and molarity for RNA libraries used in 3’ mRNA-seq in this study.

| **Sample ID** | **Library concentration** | **Library size (bp)** | **Molarity (nM)** |
| --- | --- | --- | --- |
|  | **(ng/μl)** |  |  |
| **1F** | 2.15 | 217 | 15.3 |
| **1S** | 1.02 | 198 | 7.97 |
| **2F** | 2.15 | 245 | 13.5 |
| **2S** | 1.97 | 244 | 12.42 |
| **3F** | 4.52 | 250 | 27.87 |
| **3S** | 3.86 | 243 | 24.48 |
| **4F** | 4.22 | 239 | 27.19 |
| **4S** | 2.32 | 229 | 15.63 |
| **5F** | 3.03 | 247 | 18.92 |
| **5S** | 1.74 | 249 | 10.74 |
| **6F** | 2.5 | 230 | 16.72 |
| **6S** | 2.06 | 235 | 13.52 |

Supplementary Table 3. Linear model (*lm* in R) for effects of paternal starvation on

sperm levels in ejaculate. R^2^= 0.008

|  | Sum Sq | Df | F value | Pr(>F) |
| --- | --- | --- | --- | --- |
| Treatment | -2720.41 | -2 | -139.42 | -1.33E-15 |
| Residuals | -282.928 | -29 | NA | NA |

Supplementary Table 4. Binomial GLMM (*glmer* in R) for effects of paternal starvation on sperm motility. Marginal R^2^ = 0.759

|  | Chisq | Df | Pr(>Chisq) |
| --- | --- | --- | --- |
| (Intercept) | 442.2298 | 1 | 3.54E-98 |
| scale(Time) | 107.5933 | 1 | 3.30E-25 |
| Treatment | 10.755 | 1 | 0.00104 |
| scale(Time):Treatment | 5.935633 | 1 | 0.014838 |

Supplementary Table 5. Binomial GLMM (*glmer* in R) for effects of paternal starvation on sperm curvilinear velocity (VCL). Marginal R^2^ = 0.406

|  | Chisq | Df | Pr(>Chisq) |
| --- | --- | --- | --- |
| (Intercept) | 3640.699 | 1 | 0 |
| scale(Time) | 47.63899 | 1 | 5.12E-12 |
| Treatment | 167.2854 | 1 | 2.90E-38 |
| scale(Time):Treatment | 38.89594 | 1 | 4.47E-10 |
